# YcjW: a transcription factor that controls the emergency generator of H_2_S in *E. coli*

**DOI:** 10.1101/383885

**Authors:** Lyly Luhachack, Ilya Shamovsky, Evgeny Nudler

## Abstract

Hydrogen sulfide (H_2_S) is a ubiquitous gaseous molecule that is endogenously produced in both eukaryotes and prokaryotes. Its role as a pleiotropic signaling molecule has been well characterized in mammals^1,2^. In contrast, the physiological role of H_2_S in bacteria only recently became apparent; H_2_S acts as a cytoprotectant against antibiotics-induced stress and affect the cell’s ability to maintain redox homeostasis ^3-5^. In *E. coli*, endogenous H_2_S production is primarily dependent on 3-mercaptopyruvate sulfurtransferase (3MST), encoded by *mstA*, previously known as *sseA*^3,4^. Here, we show that cells lacking 3MST acquired a unique phenotypic suppressor mutation resulting in compensatory H2S production and tolerance to antibiotics and oxidative stress. Using whole genome sequencing, we mapped a non-synonymous single nucleotide polymorphism (SNP) to uncharacterized Laci-type transcription factor, YcjW. We identified transcriptional regulatory targets of YcjW and discovered a major target, thiosulfate sulfurtransferase PspE, as an alternative mechanism for H_2_S biosynthesis. Deletion of *pspE* was sufficient to antagonize phenotypic suppression. Our results reveal a complex interaction between cell metabolism and H_2_S production and the role, a hithero uncharacterized transcription factor, YcjW, plays in linking the two.

## Main

H_2_S can be generated in numerous pathways-both enzymatically and non-enzymatically and from various substrates beyond cysteine^6^. The main pathway by which *E. coli* generate H_2_S when grown aerobically in nutrient rich LB is via 3MST, encoded by *mstA^3,4^*. Phenotypic consequences of decreased H_2_S production include greater susceptibility to multiple classes of antibiotics and H_2_O_2_ ^3,4,7^. However, we discovered that antibiotics-sensitive strain *ΔmstA* reverted to the resistant phenotype of the isogenic parent when challenged with different antibiotics. The *ΔmstA* variant, referred to as *ΔmstA-sup* in this study, was indistinguishable from wild type in time-kill assay analysis and growth curves of cells exposed to gentamicin, nalidixic acid and carbenicillin (Fig. 1a and Supplementary Fig. 1). Furthermore, this strain also had increased tolerance to hydrogen peroxide (H_2_O_2_), compared to its still sensitive parent *ΔmstA* strain (Fig. 1b). Using both the classic lead acetate reactivity test for H_2_S detection and a fluorescent-based probe, WSP5^8^, we confirmed that this phenotypic reversion was concurrent with increased H_2_S production, comparable to wild type (Fig. 1c). In contrast, significant levels of H_2_S remained undetectable in *ΔmstA* till OD_600_ 1.5.

**Figure 1.**
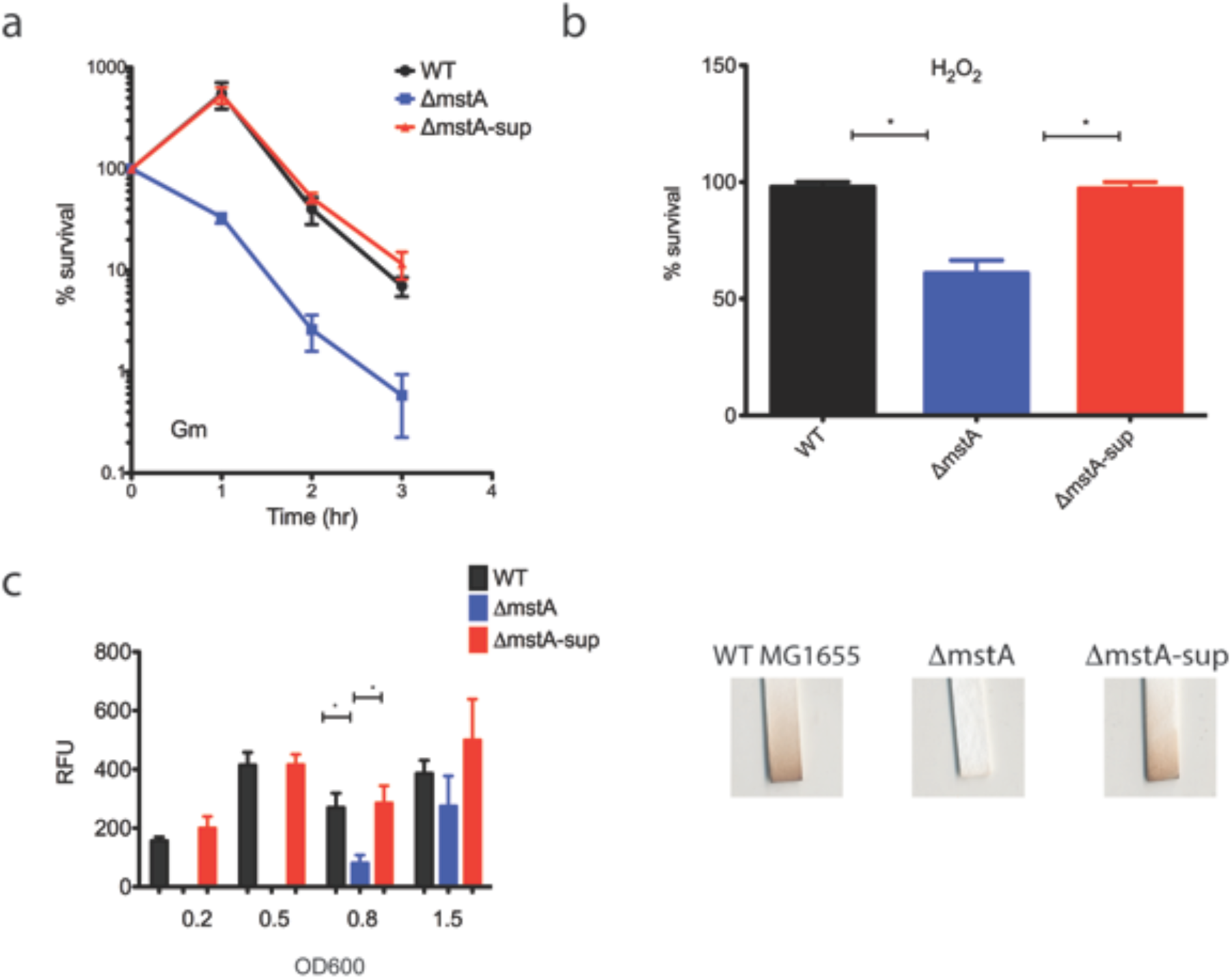
*E. coli* MG1655 lacking 3MSTA acquires phenotypic suppression and has increased H_2_S levels and tolerance to Gm and H_2_O_2_. (a) *ΔmstA-sup* has increased survival rate compared to *ΔmstA* when treated with 2ug ml^−1^ gentamicin in a time-kill assay (b) *ΔmstA-sup* also has increased tolerance after exposure to 5mM H_2_O_2_ for 30 minutes (c) H_2_S production as measured with fluorescent probe, WSP5. Relative fluorescent units are normalized to OD**600** and minus the background fluorescent of PBS buffer+100uM L-cysteine and WSP5. H_2_S reacts with lead acetate, leading to staining of strips (Sigma-Aldrich). Values are means±SD (n=3). * p<0.05

We utilized whole genome sequencing to identify possible SNPs coding regions that could be responsible for the observed phenotypic suppression. We mapped and validated by PCR a single missense mutation unique to ΔmstA-sup to uncharacterized transcription factor, *ycjW*. The nucleotide substitution, G to A on the coding strand, results in an amino acid change from serine to asparagine at residue 258 (Fig. 2a).

**Figure 2.**
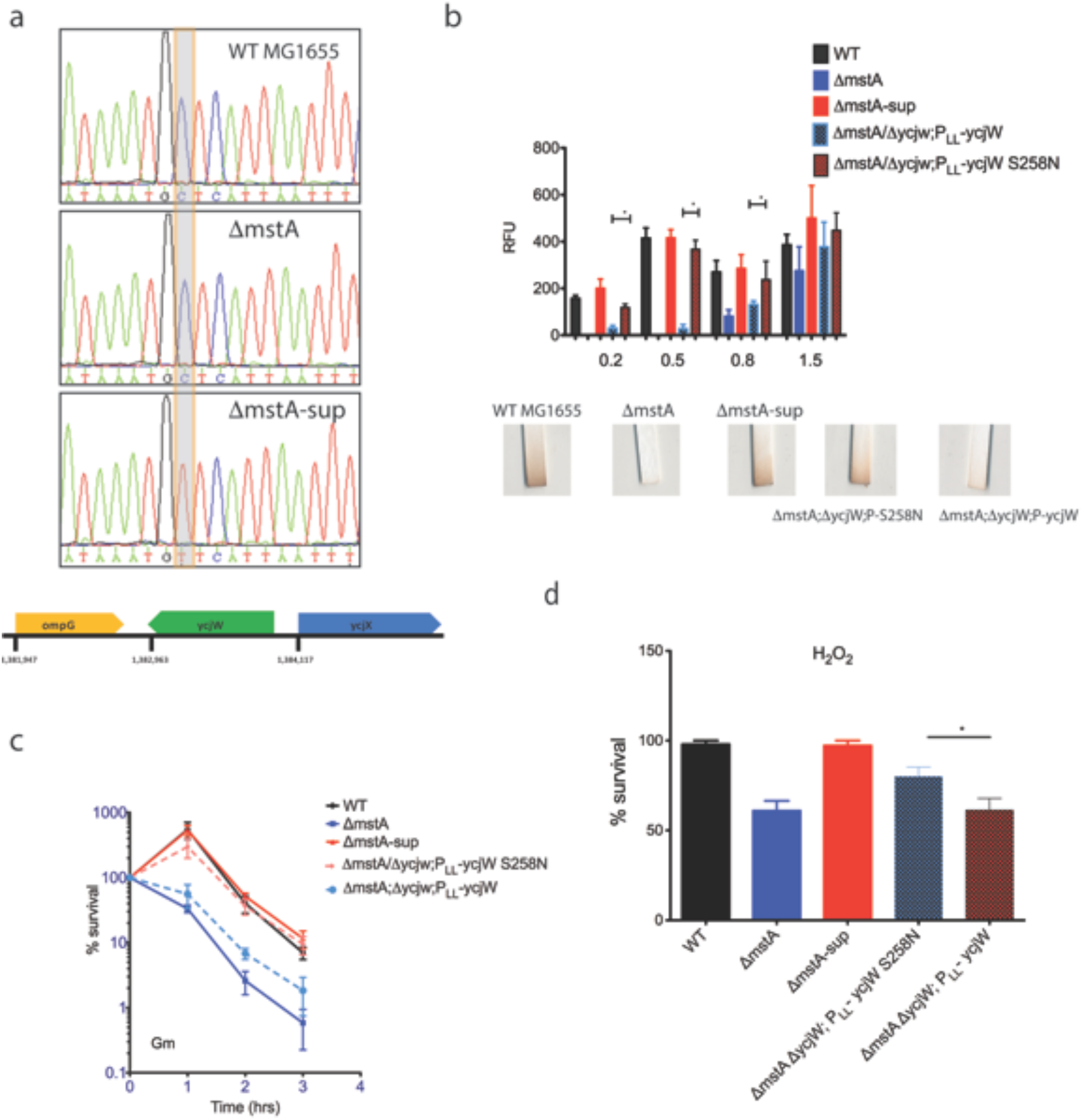
Unique nonsynonymous SNP in A *mstA-sup* is mapped to putative transcription factor *ycjW*. (a) PCR validation of whole genome sequencing. Displayed are sequences from E. coli MG155, *ΔmstA*, and *ΔmstA-sup* of (-) strand. The SNP changes amino acid 258 from serine to asparagine. (b) *ΔmstA/ΔycW* strains with plasmid expressed P_LL_-ycjW or P_LL_-ycjW S258N were measured for H_2_S production. Only *ΔmstA/ΔycW;* P_LL_-ycjW S258N had increased H_2_S production with no significant differences with WT or *ΔmstA-sup*. At OD_600_ 1.5 no significant differences exist between strains. (c) Cells exposed to 2ug ml^−1^ Gm. *ΔmstA;ΔycW;* P_LL_-ycjW S258N had increased survival rate. *ΔmstA/ΔycW;* P_LL_-ycjW was more sensitive to Gm compared to wild type and *ΔmstA-sup*. (d) *ΔmstA/ΔycW;* P_LL_-ycjW S258N improved tolerance to H_2_O_2_. Values are means±SD (n=3) for all experiments.

YcjW is annotated as a putative member of the LacI/GalR family of repressors that are largely responsible for carbohydrate metabolism. Common features of the family include an N-terminal helix-turn-helix DNA-binding domain, a linker domain, and a C-terminal ligand-binding domain ^9^. To investigate SNP functionality, we constructed two strains, bearing a plasmid expressing either wild type YcjW (pLLY1) or S258N YcjW (pLLSN1), in the background of *ΔmstA/ΔycjW*. Figure 2b shows that only plasmid-expressed mutated YcjW is able to restore H_2_S production, quantitated by utilizing the WSP5 probe and qualitatively shown by lead acetate assay. Furthermore, only *ΔmstA/*ΔycjW;P_LL_-ycjW (S258N) have an increased survival rate when challenged with gentamicin, H_2_O_2_ and nalidixic acid (Fig. 2c and 2d and Supplementary Fig. 2). Thus, we confirm that S258N YcjW in *ΔmstA-sup* is responsible for the increased hydrogen sulfide production and antibiotics and oxidative stress tolerance relative to *ΔmstA*.

To identify transcriptional targets of YcjW, we performed ChIP-seq using an antibody against chromosomal 3×FLAG-tagged YcjW from wild type, and *ΔmstA* cells, and 3xFLAG-tagged YcjW S258N from ΔmstA-sup cells. Figure 3a shows representative peaks identified by MACS2^10^ from aligned sequence reads. The most enriched regions, for all three strains, are at two sites near *ycjW;* the first site is before the translation start site of *ycjM* but after a predicted transcription start site and the second lies between *ycjT* and *ycjU*. The binding motifs for LacI type family transcription factors are typically palindromes with a conserved central CG pair ^11^. Recently, Zuo and Stormo experimentally tested the predicted binding motif for YcjW ^12^. Combined with our analysis of peak summits, we found the same sequence in our data. Using the putative binding sequence, we further restricted peaks to ones containing the conserved 14bp motif, allowing up to three mismatches and with a fold enrichment greater than 5. With those criteria, we identified two additional peaks specifically in *ΔmstA*, that are not enriched in wild type or *ΔmstA-sup*. The two sites are near the promoter of *narP*, encoding a two-component nitrate/nitrite response system and the other is located within the coding sequence of *cyaA*, encoding adenylate cyclase (Supplementary Fig. 3).

**Figure 3.**
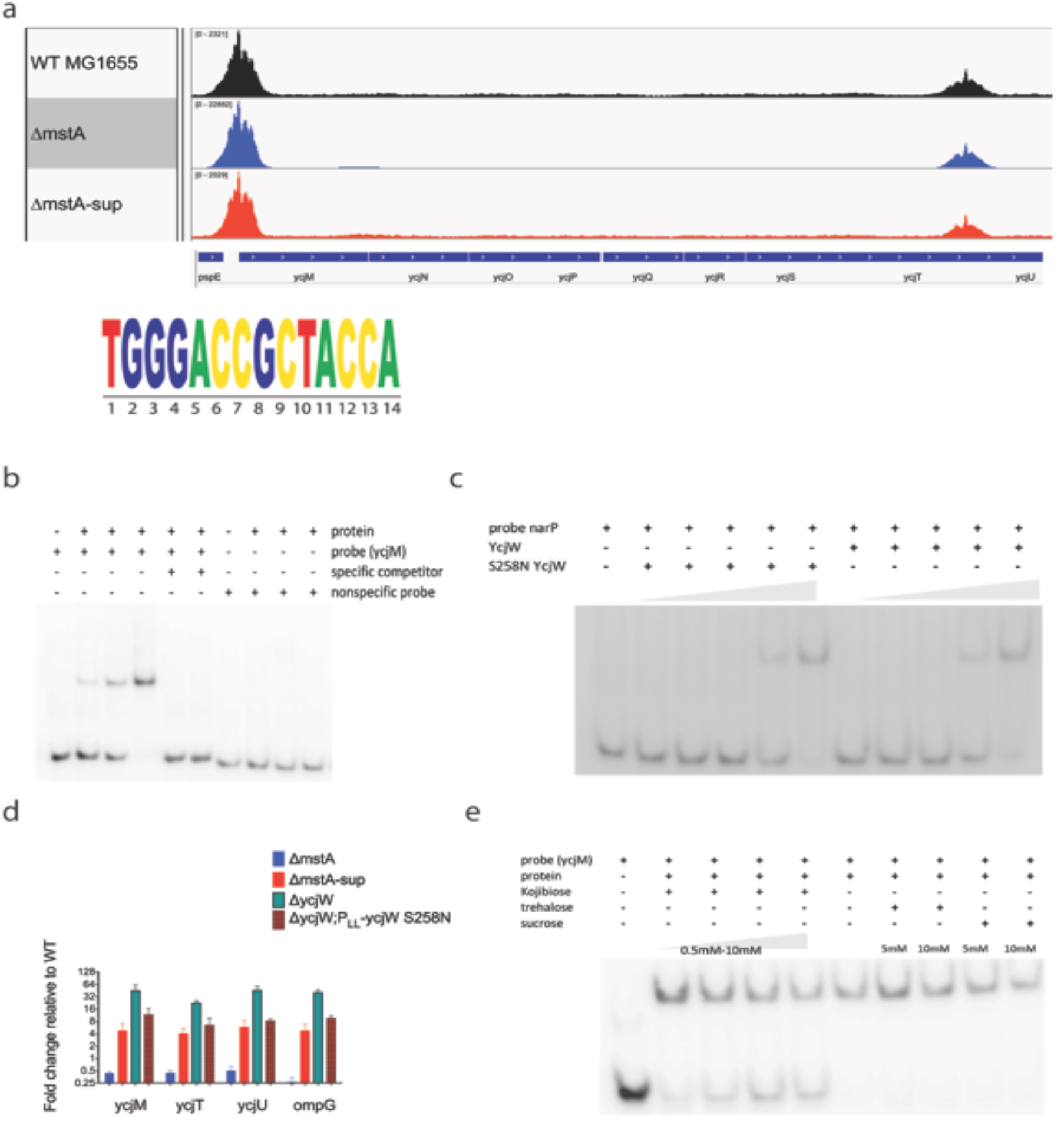
YcjW, shows binding enrichment near *ycjM* and ycjU and regulates expression of operon *ycjMNOPQRSTUV-ompG*. (a) Represented on Integrative Genomics Viewer (IGV), are sorted, aligned sequences containing pileup data to reference genome NC_000913.3. MACS2^10^ was used for peak calling. Enriched peaks are upstream of *ycjM* and *ycjU*. A 14 nucleootide sequence identified as the binding motif for YcjW. (b) YcjW protein was titrated to DNA:protein ratios of 1:0.5, 1:1, and 1:2. Unlabeled ycjM probe was added to the reaction in excess to compete for binding. (c) YcjW and S258N YcjW protein were titrated to DNA:protein ratios of 1:0.125, 1:0.25, 1:0.5, 1:1, and 1:2. with *narP* probe. (d) qRT-PCR of a subset of genes in the *ycjM-V* and *ompG* operon. The absence of *ycjW* results in massive upregulation. *ΔmstA-sup* and *ΔycjW;P_LL__ycjW* both showed moderate and significant increased expression while mRNA levels are repressed in *ΔmstA*. Values are means±SD (n=3). (e) YcjW protein was pre-incubated with Kojibiose, trehalose, or sucrose before radiolabeld DNA probes were added to the mixture. Only kojibiose prevented complex formation at a concentration of 0.5mM. In contrast, both sucrose and trehalose added in excess at 5mM and 10mM failed to disrupt binding.

We then validated transcription factor binding through electrophoretic mobility shift assay (EMSA). We designed 50bp DNA probes containing the predicted binding sequence in the center. The YcjW protein reduced the mobility of the upstream *ycjM* DNA probe at about a 1:0.5 DNA:protein ratio. Increasing amounts of protein corresponded to an increase in YcjW-DNA complex (Fig. 3b). YcjW (S258N) also reduced DNA probe mobility at the same DNA:protein ratio, using *narP* probe (Fig. 3c). Titration of the normal protein and S258N YcjW showed that they both bound DNA probe starting at a DNA:protein ratio of 1:0.5. At a ratio of 1:2, no free DNA probe could be detected.

The region downstream of *ycjM* contains a predicted operon consisting of 10 *genes-ycjMNOPQRSTUV* and *ompG*. To test functionality of transcription factor-DNA binding to gene expression, we determined the amount of relative mRNA fold change using qRT-PCR. In Δ*ycjW*, representative genes, *ycjM, ycjT, ycjU*, and *ompG* are significantly upregulated, confirming that YcjW is a repressor (Fig. 3d). The absence of YcjW results in constitutive derepression of its regulatory transcriptional targets. The same genes exhibit a similar pattern, increased expression, in *ΔycjW;P_LL_-ycjW* S258N relative to wild type MG1655 but not to the extent of its isogenic parent, *ΔycjW*. Consistent with *ΔycjW;P_LL_-ycjW* S258N, those genes are also upregulated in ΔmstA-sup, but not *ΔmstA*, suggesting that S258N YcjW affects DNA occupancy *in vivo* but not necessarily *in vitro*. Mutational analyses of other LacI-TFs have demonstrated that a single amino acid change in the C-terminal can alter effector or co-repressor binding and therefore DNA affinity at target sites ^13,14^. The SNP is located in the C-terminal effector pocket of the protein, thus raising the possibility that it broadens specificity of inducer recognition, co-repressor binding affinity or oligomerimerization.

qRT-PCR of *narP* and *cyaA* showed no significant change in *ΔycjW*. A small subset of genes regulated by NarP was tested as downstream targets (Supplementary Fig. 4). Two genes, *nrfA* and *ydhU* did have a modest increase while two others didn’t. NarP regulation however is complex and involves multiple regulators. Therefore, it is difficult to assess if YcjW-binding upstream of *narP* and *cyaA* is functional.

Because many Laci-type repressors act locally in response to some specific effector, we sought to identify the inducer for YcjW by considering its targets. YcjT is homologous to kojibiose phosphorylase from *Thermoanerobacter brockii* and *Pyrococcus* sp. Strain ST04 ^15,16^. Kojibiose phosphorylase can reversibly catabolize kojibiose to D-glucose and beta-D-glucose 1 phosphate. The downstream gene, *ycjU*, has experimentally been shown to encode a beta-phosphoglucomutase ^17^. Again, utilizing EMSA, we tested to see if kojibiose is the effector molecule for YcjW. The addition of kojibiose at 1mM disrupts the YcjW-DNA complex (Fig. 3e). Other disaccharides tested in excess of up to ten times, trehalose and sucrose, did not affect binding. However, attempts to grow *E. coli* K-12 MG1655 on minimal media with kojibiose as the sole carbon source were unsuccessful (data not shown and ^18^). Growth on EZ Rich Defined media supplemented with kojibiose as the carbon source did grow but had a rather pronounced defect. Deletion of *ycjW* did not improve growth rates either (Supplementary Fig. 5). However, the concentration of kojibiose added to media is limited by its low solubility. It is possible that a higher concentration of kojibiose supplied would support enhanced growth. Taken together, our results indicate that kojibiose might not be the natural inducer of YcjW but perhaps some derivative of kojibiose. Recently, the substrate for YcjM was identified as glucosylglycerate, alongside kojibiose for YcjT ^18^. Glucosylglycerate is an osmoprotectant in bacteria and archaea and accumulates under salt stress and limited nitrogen availability ^19,20^. However, most of our experiments were conducted in LB with amino acids constituting the main carbon source. We find it unlikely that synthesis of either the glycoside or disaccharide could occur without the appropriate substrate, and therefore is not likely involved in *ΔmstA* phenotypic suppression.

While it is not directly evident how a cluster of carbohydrate catabolic genes regulated by YcjW can lead to an alternative pathway for H_2_S production, *pspE* encoding a thiosulfate sulfurtransferase, lies immediately upstream of *ycjM*. PspE has mercaptopyruvate sulfurtransferase activity, albeit low compared to thiosulfate ^21^. It is part of a cluster of genes known as the phage shock operon, consisting of *pspABCDE*. Although it can be co-regulated with the other *psp* genes, it can also be transcribed independently from its own promoter^22^. PspF is the transcriptional activator and phage shock protein A, encoded by *pspA*, negatively regulates PspF^23^.

*pspE* mRNA expression is significantly increased in ΔmstA-sup and *ΔycjW* relative to wild type cells. In contrast, there are no significant differences in relative expression of the other two thiosulfurtransferase genes (Fig. 4a). Furthermore H_2_S production is undetectable during early exponential and mid-log phase in *ΔmstA-sup/ΔpspE* (Fig. 4b). However, at late logarithmic phase, H_2_S levels are now detectable to the same degree as MG1655 and ΔmstA-sup. In addition, overnight incubation with lead acetate strips shows no discernable difference in H_2_S extracellular production between the three strains (Supplementary Fig. 6). We conclude from the significant delay of H_2_S generation in *ΔmstA-sup / ΔpspE* that PspE is capable of generating H_2_S in early growth phases as observed in *ΔmstA-sup*. However, at later growth stages, another pathway for H_2_S production is activated and/or PspE is no longer sufficient. Moreover, *ΔmstA-sup/ΔpspE* also has increased sensitivity to gentamicin treatment compared to *ΔmstA-sup* and *E. coli* MG1655 but not quite as sensitive as *ΔmstA*. Overexpression of PspE in *ΔmstA* increases survival rate but only to the extent of *ΔmstA-sup/ΔpspE*, not *ΔmstA-sup* or wild type (Fig. 4c). Altogether, we conclude that the SNP in *ycjW* resulted in increased expression of *pspE* in ΔmstA-sup. This is sufficient but not wholly responsible for increased H_2_S biosynthesis and in turn, the phenotypic suppression observed in ΔmstA-sup. We propse a model wherein, *E.coli* cells lacking MstA acquires a SNP in transcription factor YcjW. The SNP imparts moderate constitutive expression of YcjW targets, one of which is *pspE*. Thiosulfate sulfurtransferase PspE is then able to increase H_2_S production in *ΔmstA*, and subsequently protect the cells from antibiotics and H_2_O_2_ induced stress (Fig. 4d).

**Figure 4.**
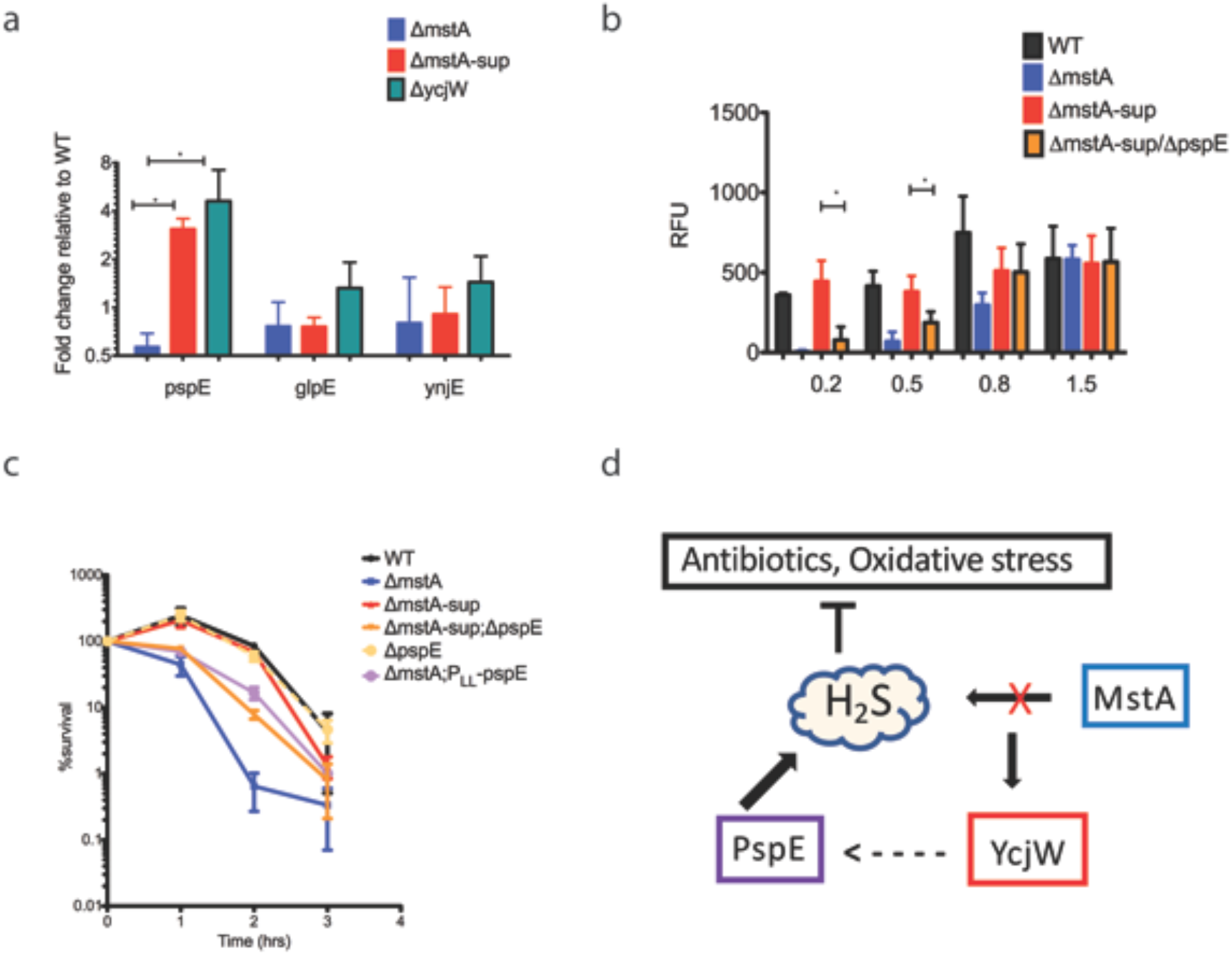
Deletion of *pspE* in A *mstA-sup* decreases H_2_S biosynthesis during exponential growth. (a)qRT-PCR of three thiosulfate sulfurtransferase genes. RNA was isolated from cells grown to OD_600_ ~ 0.4. Only *pspE* expression was significantly increased in both *ΔmstA-sup* and *ΔycjW* (b) H_2_S levels in *ΔmstA-sup; ΔpspE* were undetectable till OD_600_ reached 0.8. (c) *ΔmstA-* sup/ΔpspE had increased sensitivity to gentamicin compared to WT and *ΔmstA-sup*. Overexpression of pspE in *ΔmstA* increased tolerance to gentamicin comared to *ΔmstA*. Values are means±SD (n=3) (d) In cells lacking MstA which is responsible for H_2_S production, S258N YcjW upregulates PspE as an alternate route of H_2_S biosynthesis.?

The region upstream of *pspE* does not contain a strong binding motif for YcjW, nor do the regions flanking regulators PspA and PspF. This is not entirely unexpected since none of the other genes tested in the *psp* operon were upregulated in *ΔmstA-sup* or *ΔycjW* (Supplementary Fig. 6). Additionally, the moderate but significant increase of *pspE* mRNA in *ΔycjW*, in comparison to *ycjMNOPQRSTUV-ompG*, suggests a more complicated interaction than direct DNA-binding. It may be indicative of “leaky” expression driven by the proximity of the 3’end of *pspE* to predicted transcription start sites for *ycjM* and the binding site of YcjW. Perhaps, YcjW acts via a co-repressor. Another possibility is that the metabolic state of *ΔmstA-sup* with mutated *ycjW* induces *pspE* as a consequence. And finally, the enzyme(s) that is induced in *ΔmstA-sup; ΔpspE*, after exponential growth, has not been identified and remains a target for future studies.

Altogether, our results reveal an alternative H2S generator (PspE) that is mobilized as a result of rapid evolutional adaptation to antibacterial stress. The SNP in YcjW, regulating metabolism to at least two rare sugars, presents an interesting link between sulfur metabolism and carbon availability. Moreover, a SNP in YcjW reflects the striking genetic plasticity employed by bacteria to promptly adapt to environmental changes and stimuli and highlights the survival advantage imparted by endogenous H_2_S.

## Acknowledgments

This work was supported by the NIH grant R01 GM126891, Blavatnik Family Foundation, and the Howard Hughes Medical Institute.

## Methods

### General growth conditions

For the general cultivation of *E.coli*, strains were grown in LB broth supplemented with kanamycin (50ug ml^−1^), or chloramphenicol (30ug ml^−1^) as appropriate. Growth on solid medium contained 1.5% agar added to LB. Where noted, MOPS EZ Rich Defined Medium Kit (Teknova) was used in place of LB^24^. Cysteine solutions were prepared immediately before use, as needed.

### Construction of strains and plasmids

For a list of all strains used throughout this work, refer to Table 1.1. BW25112 and its derivatives are from the *E. coli* Keio Knockout Collection^25^ (Thermo Scientific). Introduction of new mutations into *E.coli* MG1655 were achieved through P1 transduction as previously described^26^. Temperature-sensitive FLP recombinase plasmid pCP20 was used for the excision of selective markers as needed^27^. All constructs were verified with PCR and sequencing. Primers used throughout this study are listed in Table 1.2.

**Table 1.**

| **STRAIN NAME** | **Description** | **SOURCE** |
| --- | --- | --- |
| *E. coli* K12 MG1655 | F- lambda- *ilvG*- *rfb*-50 *rph*-1 | 31 32 |
| $\Delta mstA$ | | 3 |
| $\Delta mstA$-sup | $\Delta mstA; ycjW^{S258N}$ | This study |
| $\Delta mstA; \Delta ycjW$ | | This study |
| $\Delta mstA; \Delta ycjW; P_{LL}\text{-}ycjW$ | | This study |
| $\Delta mstA; \Delta ycjW; P_{LL}ycjW^{S258N}$ | | This study |
| $\Delta ycjW; P_{LL}ycjW^{S258N}$ | | This study |
| WT-3xFlag | *E. coli* K12 MG1655 (*ycjW* fused to 3X FLAG) | This study |
| $\Delta mstA$-3XFlag | $\Delta mstA$ (*ycjW* fused to 3X FLAG) | This study |
| $\Delta mstA$-sup-3XFlag | $\Delta mstA; ycjW^{S258N}$ (*ycjW*<sup>S258N</sup> fused to 3X FLAG) | This study |
| $\Delta pspE$ | | This study |
| $\Delta mstA$-sup; $\Delta pspE$ | $\Delta mstA; ycjW^{S258N}; ; \Delta pspE$ | This study |
| $\Delta mstA; P_{LL}\text{-}pspE$ | $\Delta mstA$; overexpression PspE | This study |

**Table 2.**
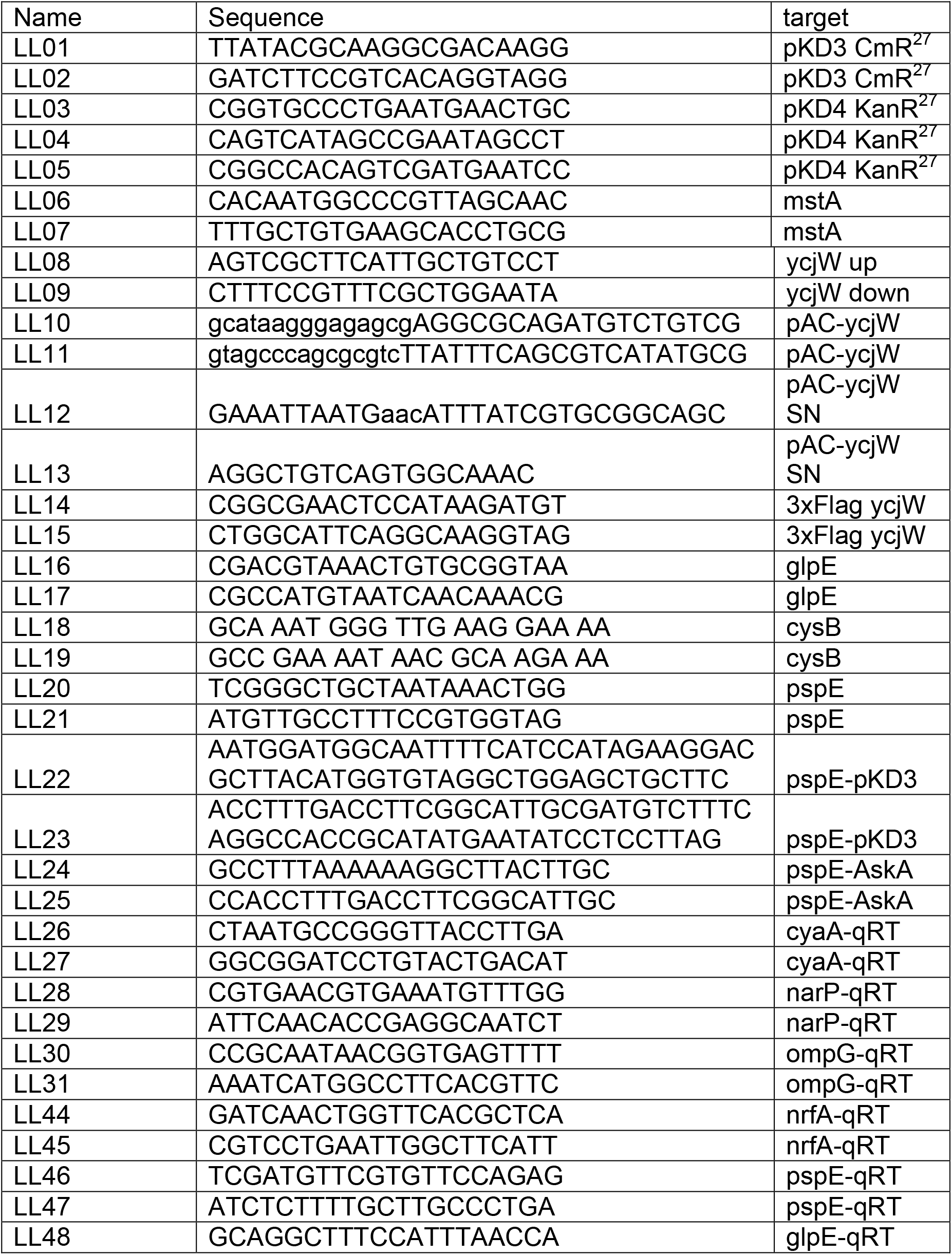
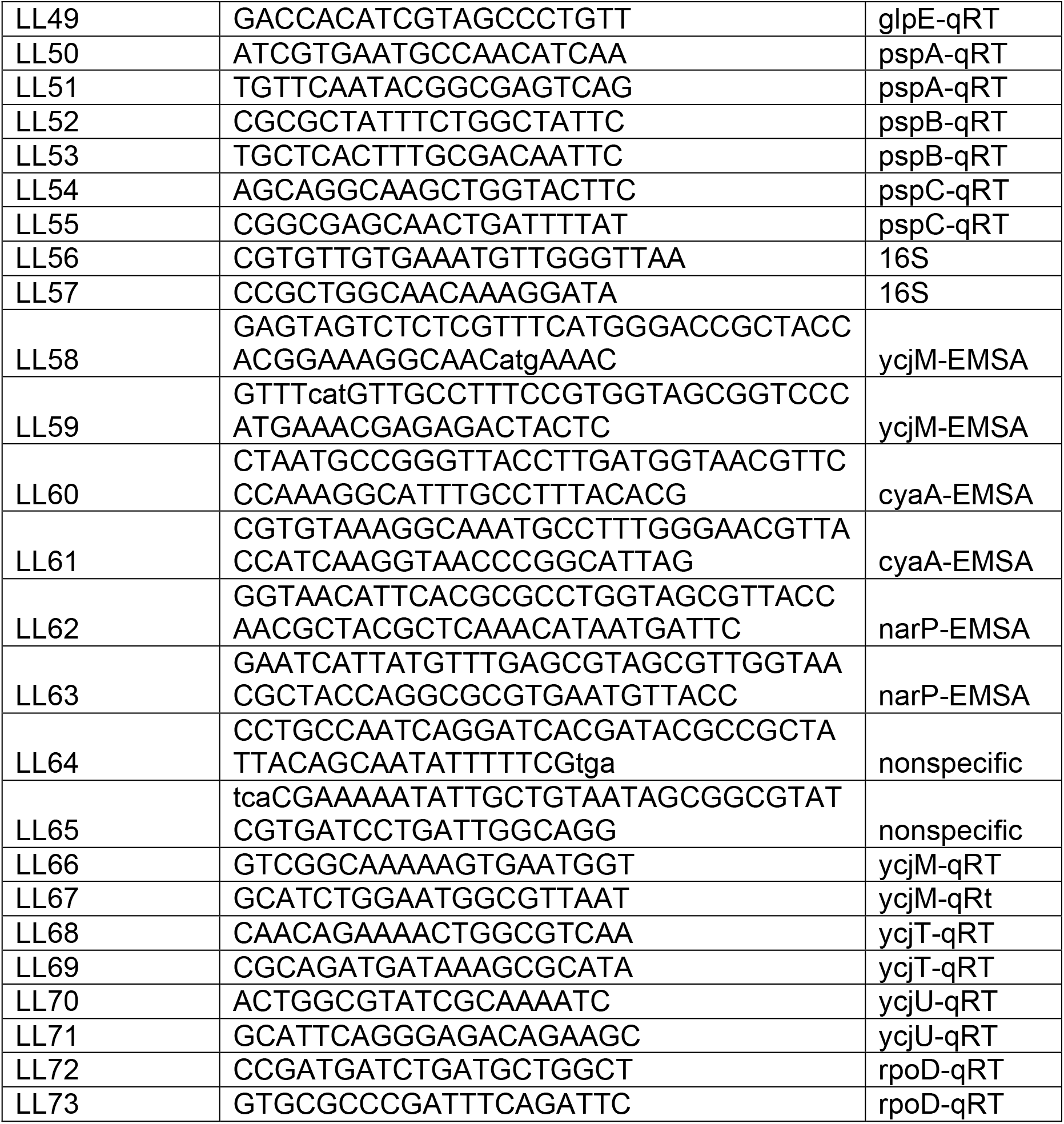

To generate pLLY1, ycjW was PCR amplified from *E.coli* MG1655 using primers LL10 and LL11 and cloned into pACYC184 plasmid (NEB) using the Gibson Assembly Mastermix, according to the manufacturer’s protocol (NEB). Plasmid pLLSN3 was generated as above except ycjW was PCR amplified from mstA-sup. The Q5 Site-Directed Mutagenesis Kit (NEB) was used to generate pLLSN1 from pLLY1, according to manufacturer’s protocol.

Transformations were performed using the CaCl_2_ competent cell protocol^28^. All plasmids were sequenced for verification.

Addition of 3xFLAG tag to *ycjW* at its chromosomal locus was achieved as previously described, with slight modifications^29^. Briefly, primers pLL14 and pLL15 were used to PCR amplify Cm^R^ cassette from pKD4. PCR product was transformed into appropriate electrocompetent strains.

### H_2_S detection

End-point detection of H_2_S production by lead acetate strips were performed as previously described^3^. Test strips were purchased from Sigma-Aldrich. Monitoring H_2_S generation with the WSP5 fluorescent probe followed a modified protocol from Peng et al.^8^ Briefly, cells were grown in LB at 37°C to desired OD_600_ and aliquots of approximately 4×10^8^ cells were taken. The extinction coefficient used for calculations is OD_600_ of 1.0 is equal to 8×10^8^ cells. A working solution of WSP5 was made immediately before use and added to cells for a final concentration of 10uM. Samples were incubated at 37°C for 30 minutes and then washed in PBS buffer, pH 7.4, to remove excess probe. Cells were resuspended in PBS buffer and incubated at room temperature for 30 minutes. Cytation3 (Biotek) was used to take fluorescent readings, at excitation 500 nm and emission 533 nm. All experiments were repeated for a total of three times. Background values were subtracted during analysis.

### Time-kill assay and growth curves

Overnight cultures of *E. coli* were diluted 1:300 into fresh media and grown to an OD_600_ of ~0.2. A 1ml aliquot was serially diluted and plated onto LB agar plates to determine initial colony forming units per ml (cfu ml^−1^) after overnight incubation at 37°C. Antibiotics were added to the cultures at indicated concentrations. Aliquots of 1ml were collected at specified time intervals, serially diluted and plated. Results from three independent experiments were plotted in GraphPad version 5.0.

Growth curves were generated from Bioscreen C automated growth analysis system as previously described^3^. Antibiotics were purchased from Sigma-Aldrich or Gold Biotechnology.

### Whole genome sequencing

Overnight cultures of *E. coli* cells were used for genomic DNA isolation. The MasterPure Complete DNA Purification Kit (Epicentre) was used to purify DNA according to the manufacturer’s protocol. DNA samples were quantified using the Quant-IT PicoGreen dsDNA assay kit (Thermo Fisher) according to manufacturer’s protocol. DNA was sheared to appropriate size with Covaris, followed by adaptor ligation. Sequencing was performed at New York University School of Medicine’s Genome Technology Center.

### Quantitative RT-PCR

Cells were grown until appropriate OD_600_ and aliquots were collected and treated with RNAprotect Bacteria Reagent (Qiagen). After 5 minutes, cells were harvested and resuspended in lysis buffer (RNase-free TE buffer, 10 mg ml^−1^ lysozyme, 100 ug ml^−1^). Trizol LS (Thermo Scientific) was used according to manufacturer’s protocol to extract total RNA. Samples were treated with DNase (Invitrogen) and purified using spin columns (Zymo Research). Superscript III reverse transcriptase (Invitrogen) was used to synthesize cDNA. qPCR reactions were amplified using Power SYBr Green PCR Master Mix (Applied Biosystems) with appropriate primer sets and cDNA template.

### ChIP-seq

ChIP was carried out as previously described with the following modifications^30^. Briefly, cells were grown at 37°C to OD_600_ ~0.4 and a final concentration of 1% formaldehyde was added for in vivo cross-linking of nucleoprotein. A final concentration of 0.5M glycine was added to the culture to quench the reaction after a 20 minute incubation. Cells were collected by centrifugation and washing twice with 1X cold Tris-buffered saline then frozen in liquid nitrogen and stored at −80. Cells were resuspended in lysis buffer (50mM Tris [pH 7.5], 100mM NaCl, 1mM EDTA, protease inhibitor [Roche], 10 mg ml^−1^ lysozyme). After incubation at 37°C, IP buffer (50mM HEPES-KOH, 150mM NaCl, 1mM EDTA, 1% Triton X100, 0.1% sodium deoxycholate, 0.1% SDS, protease inhibitor [Roche]) was added at a 1:3 ratio. DNA was sheared using ultrasonicator Covaris M220 on a 10 seconds on and 10 seconds off cycle for a total of 50 cycles.

The supernatant was incubated with 3xFLAG antibody (Biolegend) and Dynabeads Protein G (Thermo Scientific) overnight at 4°C. Samples were then washed twice with IP buffer, once with IP buffer+500mM NaCl, once with wash buffer (10mM Tris, 250mM LiCl, 1mM EDTA, 0.5% NP-40, 0.5% sodium deoxycholate), and a final wash with TE. Immunoprecipitated complexes were eluted in elution buffer (50mM Tris, 10mM EDTA, 1% SDS) at 65°C for 20 minutes. Samples were treated with RNAse A (Qiagen), at 42°C and then uncross-linked with elution buffer+pronase for 2 hours at 42°C, followed by 6 hours at 65°C. DNA was purified using ChIP Clean and Concentrate (Zymo Research). Prior to sequencing, DNA was checked on TapeStation 2200 for appropriate size (Agilent). ChIP experiments were repeated for a total of three replicates.

For sequencing, sample libraries were prepared by using the NEBNext ChIP-seq Library (Illumina), according to manufacturer’s protocol. Samples were sequenced on NextSeq 500 (Illumina). Bowtie and MACS2 were used for aligning and peak-calling, respectively^10^.

### Electrophoretic mobility shift assay

#### Protein purification

YcjW and S258N YcjW were cloned into plasmid pet28-SUMO using the Gibson Assembly Mastermix kit, according to the manufacturer’s protocol (NEB). Auto-induction media was used for protein production (152). Cells were harvested and resuspended in lysis buffer (1M NaCl, 5mM Imidaziole, 5% Glycerol, protease inhibitor cocktail [Roche]) and sonicated. AKTA Start system was used for chromatography with HisTrapHP columns (GE Healthcare Life Sciences). Columns were washed in wash buffer (50 mM Tris-Cl [pH 8.0], 10 mM Imidazole, 5% Glycerol, 500 mM NaCl), followed by gradient elution with elution buffer (50 mM Tris-Cl [pH 8.0], 250 mM Imidazole, 5% Glycerol, 250 mM NaCl). The SUMO tag was cleaved with SUMO protease in dialysis buffer (200 mM NaCl, 50 mM Tris-Cl [pH 8.0], 5% Glycerol, 1mM DTT). Samples were applied to a HiTrap HeparinHP column (GE). Columns were washed with buffer (20mM Tris [pH 8.0]. 50mM NaCl, 5% glycerol), and eluted in elution buffer (20 mM Tris [pH 8.0], 1.5 M NaCl, 5% glycerol). The sample was concentrated to 5 mL and injected onto a Superdex 200 column with GF buffer (20 mM Tris-Cl [pH 8.0], 50 mM NaCl, 1 mM DTT).

#### EMSA

dsDNA probes containing the binding sequence were radiolabeled with gamma ^32^P rATP using T4 polynucleotide kinase (NEB). Labeled probes were purified by passage through size exclusion columns (Bio-Rad). Binding reactions were done as previously described^12^. The gel was then exposed to a phosphor screen and visualized on Storm 820 Phosphorimager (GE Healthcare). Experiments with various disaccharides were done in a similar fashion, except purified protein was incubated with appropriate sugar for 20 minutes at room temperature before addition of radiolabeled probe.

---

**Fig. S1.**
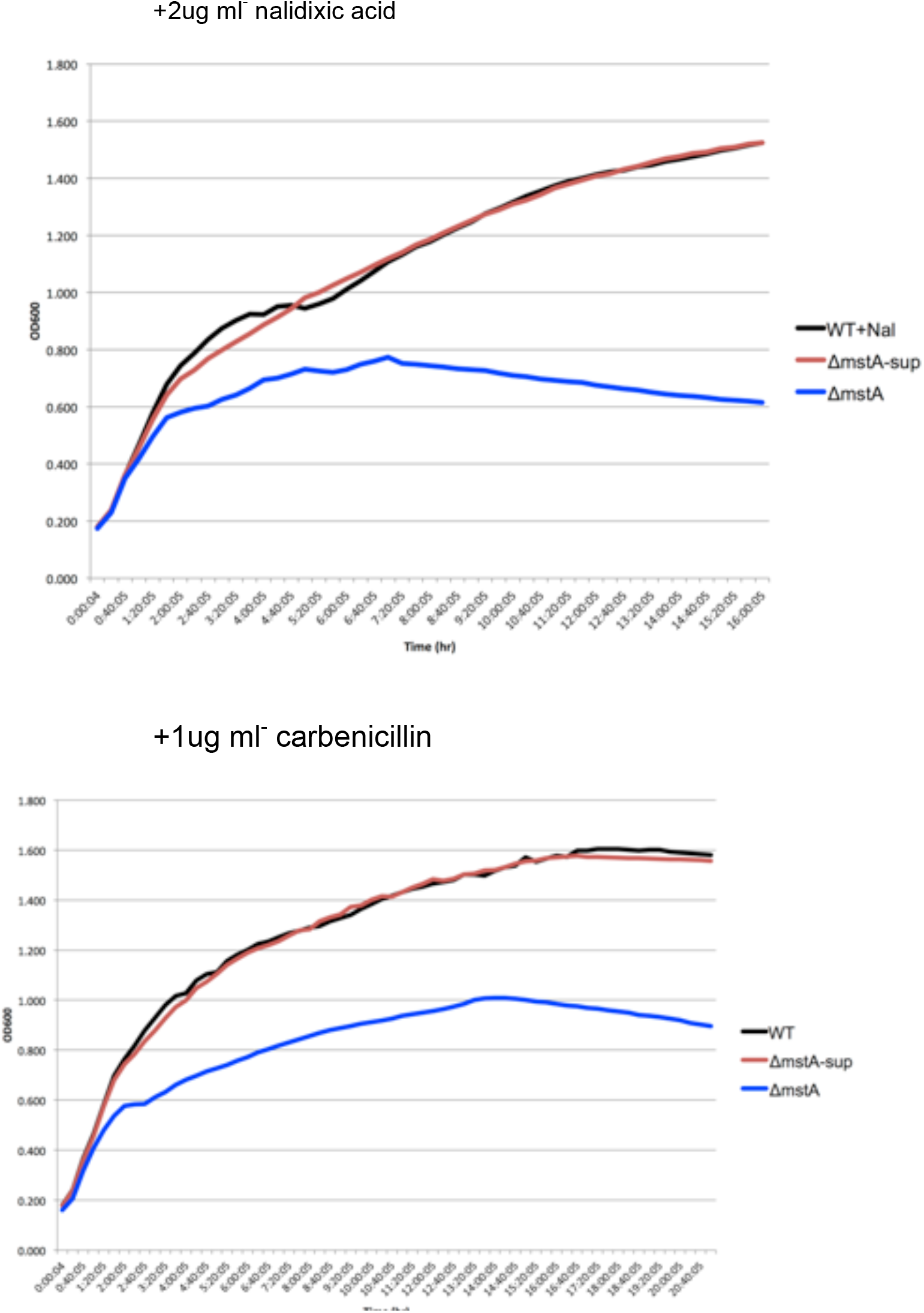
Δ*mstA*-sup has increased tolerance to carbenicillin and nalidixic acid (a) Cells were grown in the presence of 2ug ml^−1^ nalidixic acid and monitored for growth by OD_600_. *ΔmstA-sup* grows as well as wild type (b) Cells were grown in the presence of 1ug ml^−1^ carbenicillin and monitored for growth by OD_600_. *ΔmstA-sup* grows as well as wild type.

**Fig. S2.**
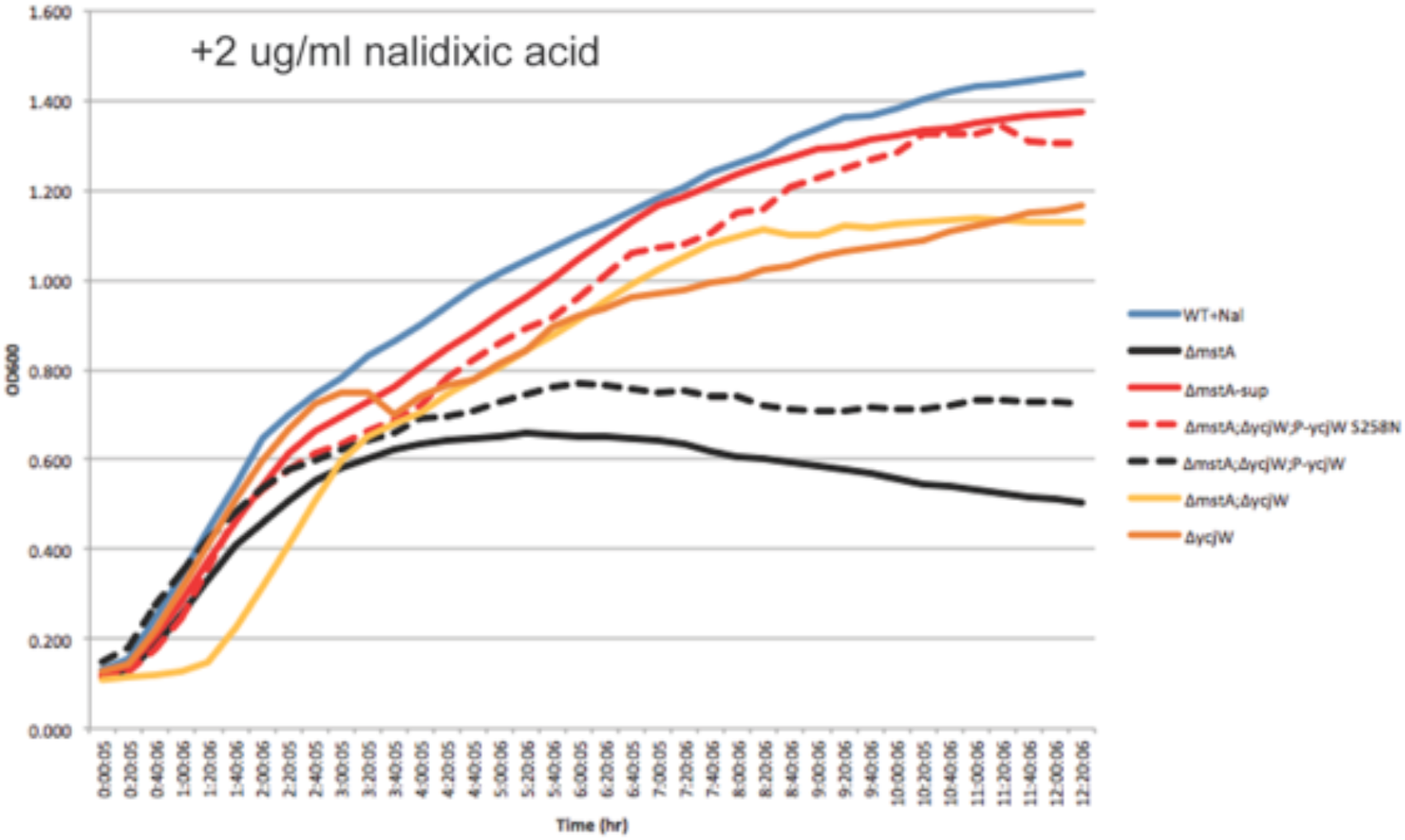
Different strains of *ΔycjW* and *ΔmstA* mutant combinations, grown in the presence of nalidixic acid. *ΔmstAIΔycW:*; P_LL_-ycjW S258N has similar growth rate to WT and *ΔmstA-sup* when grown in the presence of a sublethal concentration of nalidixic acid. In contrast, *ΔmstAIΔycW;* P_LL_-ycjW has a reduced growth rate compared to wild type and *ΔmstA-sup*. Interestingly, *ΔycjW* also has impaired growth, though not to the extent of *ΔmstA*.

**Fig. S3.**
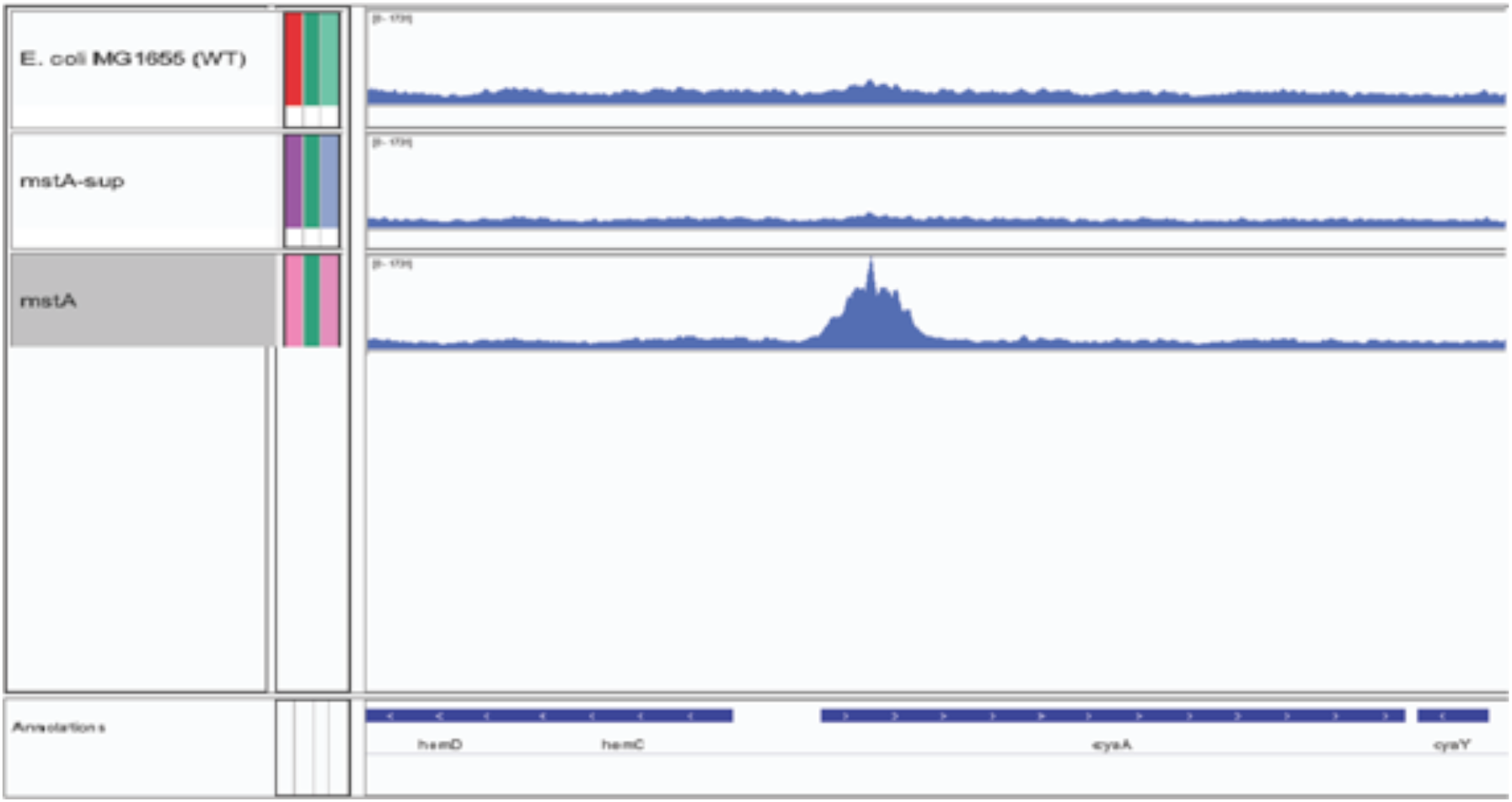
ChIP-seq aligned data and pileup data visualized on IGV. Shown is the region near *cyaA*. Only *ΔmstA* showed an enrichment of binding at a site downstream of the translation start site of *cyaA*

**Fig. S4.**
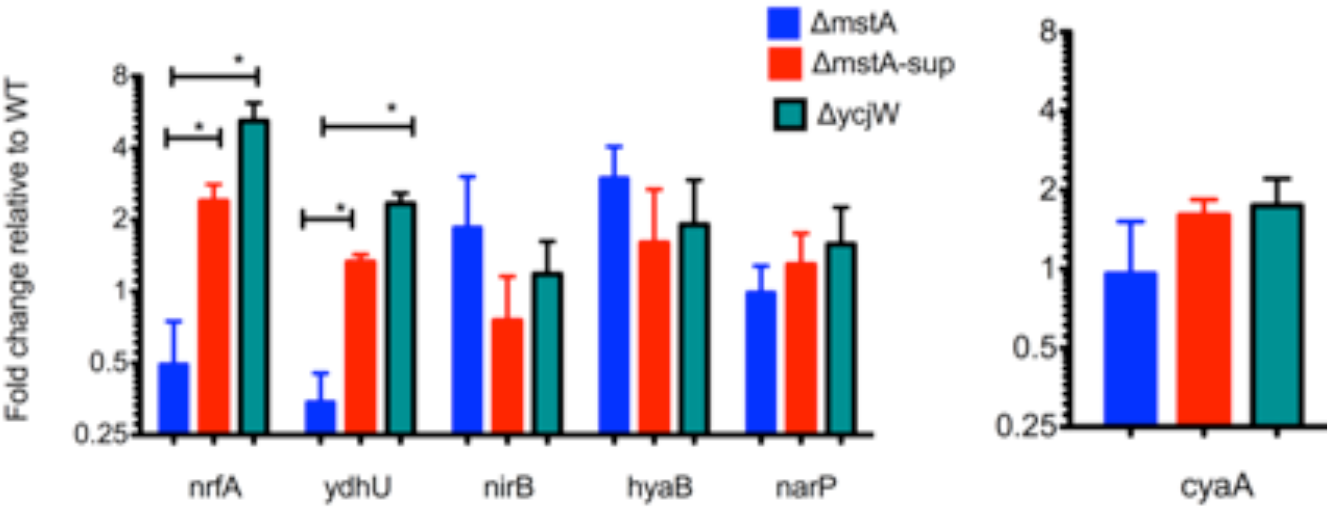
YcjW does not affect *narP* expression or *cyaA*. NarP is a two-component nitrate/nitrite response regulator and can either repress or activate transcription. Of the genes measured for changes in mRNA levels, only nrfA and ydhU are significantly increased in *ΔycW*. Upregulation however, is modest.

**Fig S5.**
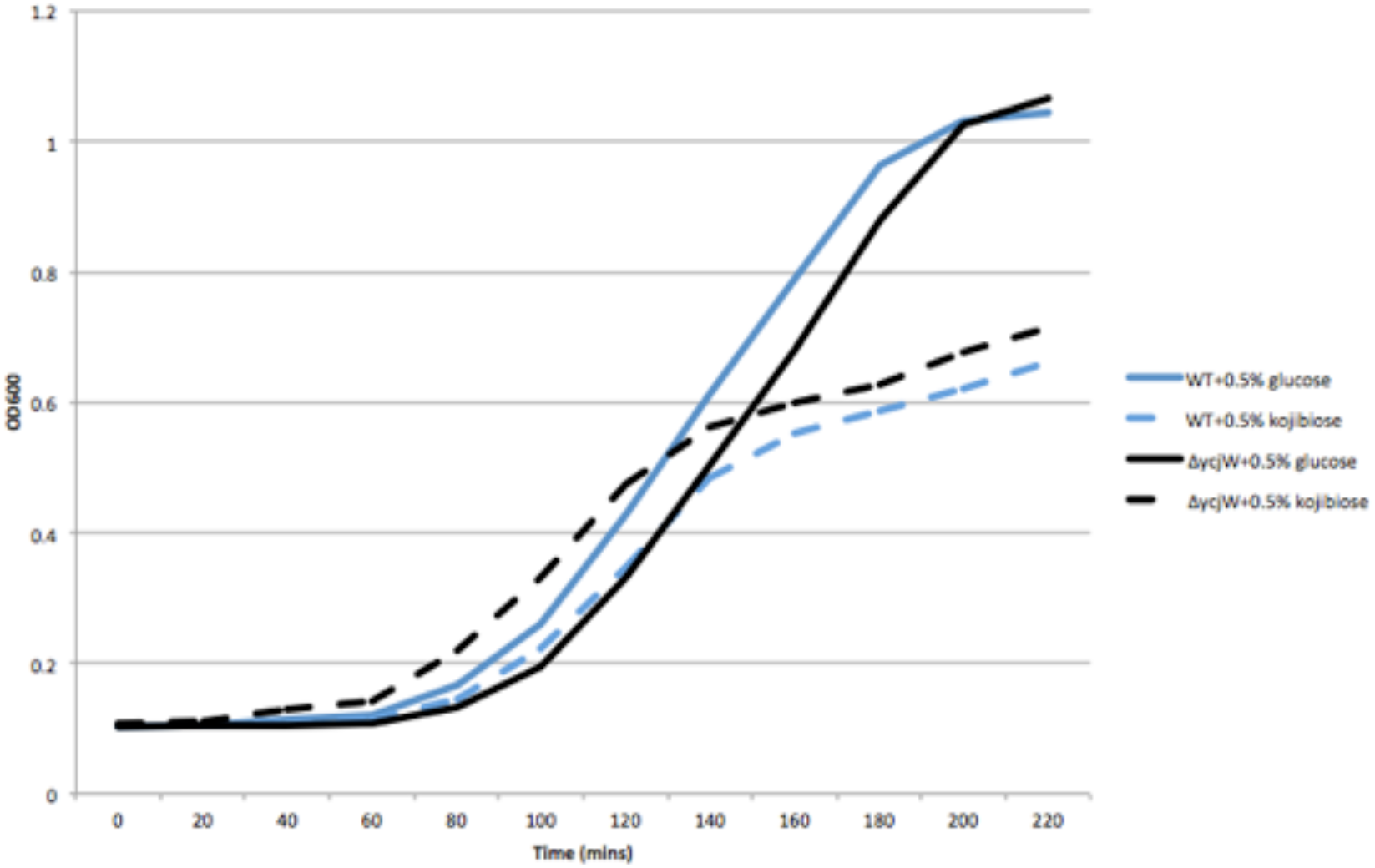
E. coli cells were grown in EZ rich defined media supplemented with either 0.5% glucose or 0.5% kojibiose as the carbon source. WT has a pronounced growth reduction with kojibiose. Knockout of *ycjW*, which results in significant upregulation of *ycjT*, capable of catalyzing kojibiose, did not improve growth.

**Fig S6.**
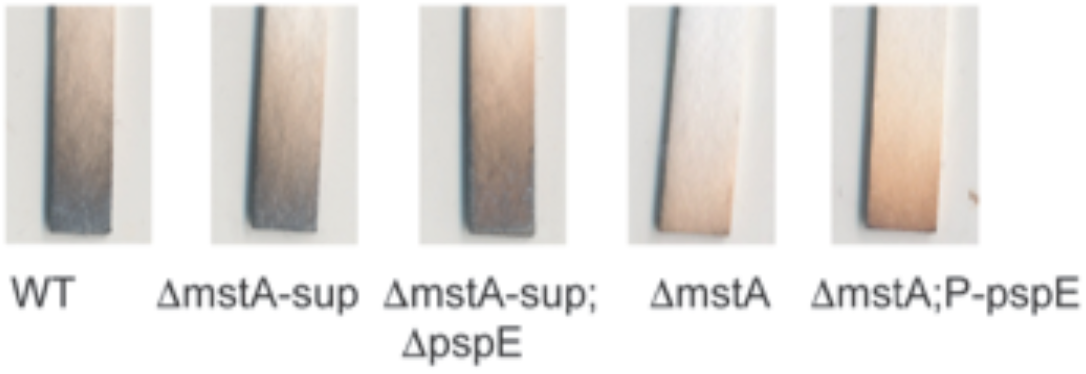
After overnight incubation with lead acetate strips, *ΔmstA-sup/ΔpspE* is able to overcome the early deficit in H_2_S production seen during exponential growth. This is in contrast to *ΔmstA*, which does not accumulate extracellular H_2_S to the extent of WTafter overnight growth.

**Fig S7.**
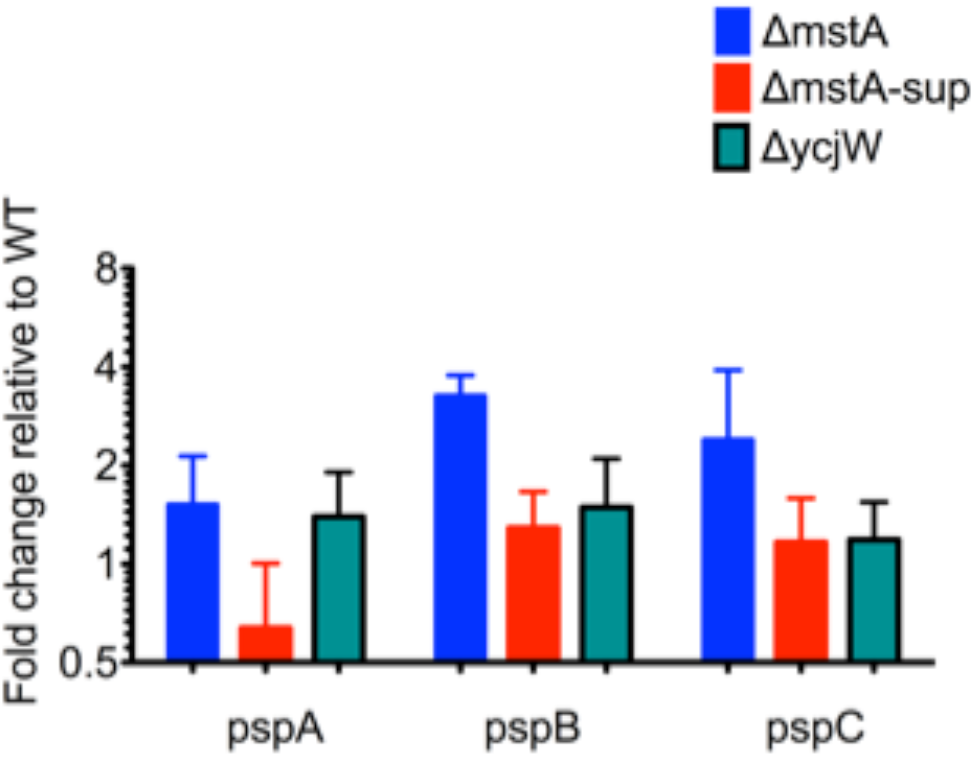
YcjW does not affect other psp genes. *pspABC* is upstream of *pspE* which is upregulated in *ΔycW* indicating that the regulation of pspE in this case is distinct and separate from the phage shock operon in general.

